# A genetic risk score to guide age-specific, personalized prostate cancer screening

**DOI:** 10.1101/089383

**Authors:** Tyler M. Seibert, Chun Chieh Fan, Yunpeng Wang, Verena Zuber, Roshan Karunamuni, J. Kellogg Parsons, Rosalind A. Eeles, Douglas F. Easton, ZSofia Kote-Jarai, Ali Amin Al Olama, Sara Benlloch Garcia, Kenneth Muir, Henrik Gronberg, Fredrik Wiklund, Markus Aly, Johanna Schleutker, Csilla Sipeky, Teuvo LJ Tammela, Børge G. Nordestgaard, Sune F. Nielsen, Maren Weischer, Rasmus Bisbjerg, M. Andreas Røder, Peter Iversen, Tim J. Key, Ruth C. Travis, David E. Neal, Jenny L. Donovan, Freddie C. Hamdy, Paul Pharoah, Nora Pashayan, Kay-Tee Khaw, Christiane Maier, Walther Vogel, Manuel Luedeke, Kathleen Herkommer, Adam S. Kibel, Cezary Cybulski, Dominika Wokolorczyk, Wojciech Kluzniak, Lisa Cannon-Albright, Hermann Brenner, Katarina Cuk, Kai-Uwe Saum, Jong Y. Park, Thomas A. Sellers, Chavdar Slavov, Radka Kaneva, Vanio Mitev, Jyotsna Batra, Judith A. Clements, Amanda Spurdle, Australian Prostate Cancer BioResource, Manuel R. Teixeira, Paula Paulo, Sofia Maia, Hardev Pandha, Agnieszka Michael, Andrzej Kierzek, David S. Karow, Ian G. Mills, Ole A. Andreassen, Anders M. Dale, The PRACTICAL consortium

## Abstract

**Background:** Prostate-specific-antigen (PSA) screening resulted in reduced prostate cancer (PCa) mortality in a large clinical trial, but due to a high false-positive rate, among other concerns, many guidelines do not endorse universal screening and instead recommend an individualized decision based on each patient’s risk. Genetic risk may provide key information to guide the decisions of whether and at what age to screen an individual man for PCa.

**Methods:** Genotype, PCa status, and age from 34,444 men of European ancestry from the PRACTICAL consortium database were analyzed to select single-nucleotide polymorphisms (SNPs) associated with prostate cancer diagnosis. These SNPs were then incorporated into a survival analysis to estimate their effects on age at PCa diagnosis. The resulting polygenic hazard score (PHS) is an assessment of individual genetic risk. The final model was validated in an independent dataset comprised of 6,417 men with screening PSA and genotype data. PHS was calculated for these men to test for prediction of PCa-free survival. PHS was also combined with age-specific PCa incidence data from the U.S. population to generate a PCa-Risk (PCaR) age that relates a given man’s risk to that of the population average. PHS and PCaR age were evaluated for prediction of positive predictive value (PPV) of PSA screening.

**Findings:** PHS calculated from 54 SNPs was very highly predictive of age at PCa diagnosis for men in the validation set (*p* =10^−53^). PPV of PSA screening varied from 0.18 to 0.52 for men with low and high genetic risk, respectively. PHS modulates PCa-free survival curves by an estimated 20 years between men in the 1^st^ or 99^th^ percentiles of genetic risk.

**Interpretation:** Polygenic hazard scores give personalized genetic risk estimates and can inform the decisions of whether and at what age to screen a man for PCa.

**Funding:** Department of Defense #W81XWH-13-1-0391

## Introduction

Prostate cancer (PCa) is a major health problem, with over one million new cases and over 300,000 prostate cancer deaths estimated worldwide in 2012^1^. An international, randomized, controlled trial showed that prostate-specific-antigen (PSA) screening resulted in a 20% reduction in PCa mortality by 20%^2^. However, due to concerns over a high rate of false positives, in addition to aggressive treatment of initially indolent disease, many clinical guidelines do not endorse universal screening and instead stress the importance of taking into account individual patient risk factors to inform the decision of whether to screen^3–5^. The goal is to avoid unnecessary screening while still identifying high-risk men for whom screening and early PCa detection can reduce morbidity and mortality.

A patient’s genetic predisposition could be critical to the decision of whether and when to offer him PCa screening. Genome-wide association studies (GWAS) have revealed genetic variants associated with increased risk of PCa^6,7^. These developments, combined with the recent accessibility of genotyping, provide an opportunity for genetic risk-informed cancer screening^8^. By combining risk information from an array of single nucleotide polymorphisms (SNPs), polygenic models can estimate an individual’s genetic risk for developing the disease^9^. It remains unclear to what extent this predicted polygenic risk could improve clinical decisions such as whom to screen for PCa and at what age.

Here we use data from 34,444 men of European ancestry from the international PRACTICAL consortium (http://practical.ccge.medschl.cam.ac.uk/) to develop a polygenic hazard score (PHS) to predict *age-related* risk of developing prostate cancer. The PHS was then tested in data from an independent study (UK ProtecT^10^) that included both genotype and PSA results, with the hypothesis that PHS would be an indicator of a patient’s inherent genetic risk for developing prostate cancer at various ages in his lifetime.

## Methods

### Participants

Discovery Set: For PHS model development, genotype and age data were obtained from 21 studies of the PRACTICAL consortium (Table 1), representing 31,747 men (18,868 cases, 12,879 controls) of genotypic European ancestry. Age was either at PCa diagnosis or last follow-up (for controls. Genotyping was performed via a custom Illumina array (iCOGS), and quality control steps were applied as described previously^6^. 201,043 SNPs were available for analysis.

**Table 1:**
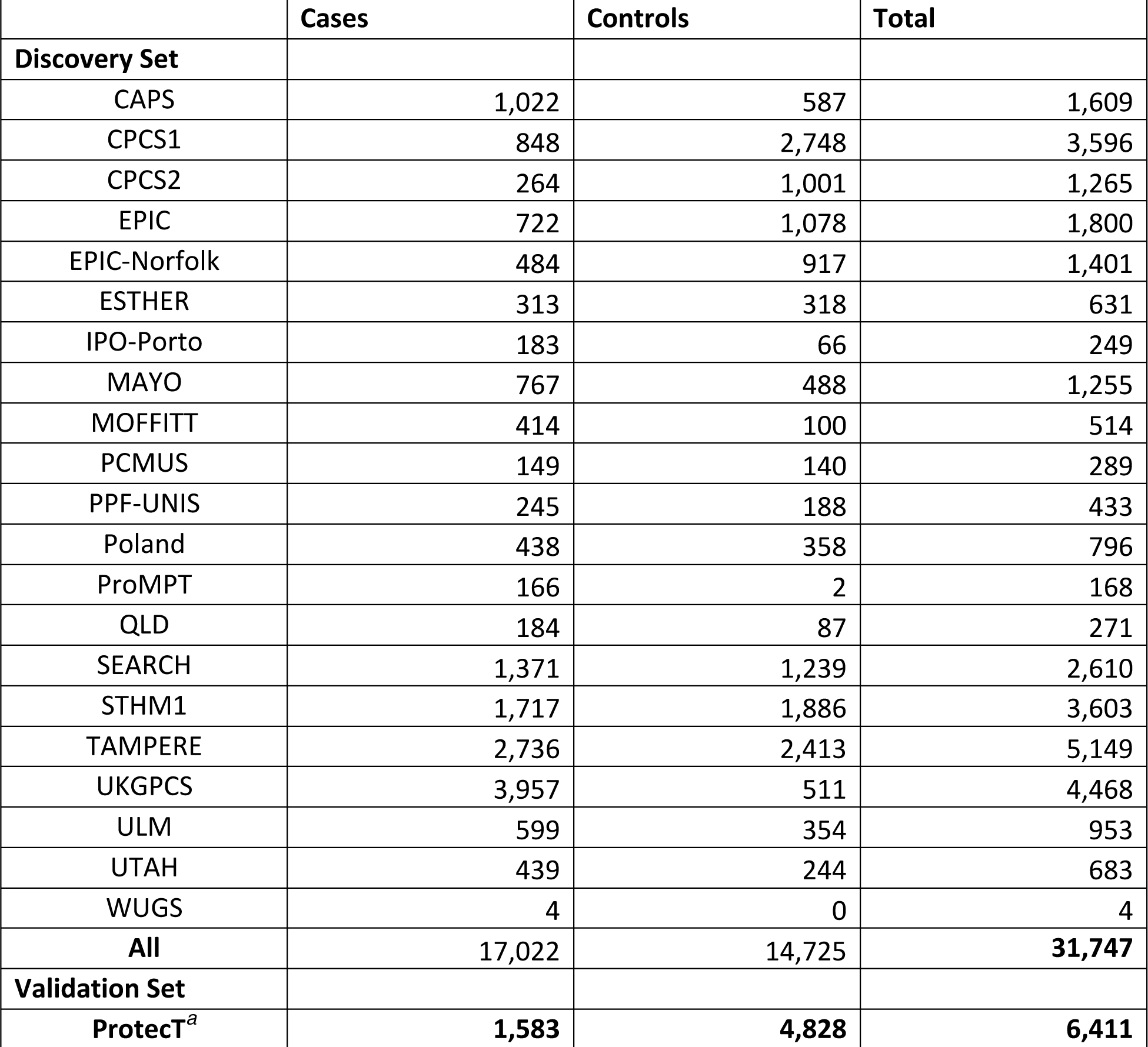
Study names and participant numbers

a Includes the 31 cases and 3,364 controls who participated in both ProtecT and UKGPCS

Validation Set: The model performance was examined in an independent study. The Validation Set comes from the ProtecT study, which screened 82,429 men with PSA testing and found 8,891 men with PSA greater than the specified threshold of 3.0 g/L or higher, of whom 2,896 were diagnosed with PCa^10^. Among those individuals, we obtained genotype and age data for 6,411 men (1,583 cases, 4,828 controls). This data set was selected for validation because PSA results were also available for all participants at time of either diagnosis or interview.

### Polygenic hazard score (PHS)

The PHS was developed previously as a parsimonious, survival-analysis model to predict the time to event outcome. It has been published elsewhere^11^. Because prostate cancer risk increases with age^12^ and anticipated age of developing prostate cancer is highly relevant to clinical management, we applied PHS for deriving both predicted absolute risk and potential age at PCa onset. In brief, a univariate trend test was applied to the entire Discovery Set (31,747 patients x 201,043 SNPs) to assess association with case or control status. All SNPs with resulting *p*-values <10^−6^ in the trend test were then entered in a forward, stepwise, greedy algorithm, to select the most predictive SNPs. In each step, logistic regression was used first to improve computational efficiency. SNPs were selected for the model only if they improved prediction of case-control status. After forward, stepwise selection, coefficients for selected SNPs were estimated using a Cox proportional hazard model to predict age at diagnosis with PCa.

The polygenic hazard score (PHS) is defined as the vector product of a patient’s genotype (*X_i_*) for the *n* selected SNPs and the corresponding parameter estimates (*β_i_*) from the Cox proportional hazards regression.

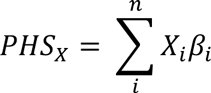

To verify whether the PHS accurately predicts age of prostate cancer onset, the PHS was calculated for all patients in the Validation Set. These values were then tested as the sole predictive variable in a Cox proportional hazards regression model for age of diagnosis. Statistical significance was set at alpha of 0.01.

### Estimate of absolute risk

The population risk of prostate cancer was estimated using methods described previously^13^. U.S. population risk data is reported by the American Cancer Society, with tables indicating the probability of developing PCa during specified age intervals, assuming the person is alive without PCa at the beginning of the interval^12^. Tables are constructed from the Surveillance, Epidemiology, and End Results (SEER) database.

Data from four such tables were used, representing twelve years of SEER data in the period from 1995 to 2012^12,14–16^. Estimates were derived of age-specific incidence as follows. For example, for the period of 2010-2012, the probability of developing PCa from age 50 to 59 was 2.1%^12^, so at a mean age of 54.5 years, we estimated the age-specific incidence as 2.1%/10 years = 0.21%/year. This was done for each reported probability in the four publications cited. Age intervals prior to age 40 were excluded due to the very low incidence in the general population. For age intervals from 70 to death, the end of the interval was taken to be 83, given an average life expectancy of 14 years for a 70-year-old man^17^. The age-specific incidence data points from all four publications were then fit to an exponential curve using linear regression in order to develop a continuous model of age-specific incidence in the U.S. SEER population.

### Examining impact of genetic risk on PSA screening

To assess the clinical significance of PCa PHS, we looked at the positive predictive value (PPV) of PSA testing within the Validation Set, with clinical diagnosis (including biopsy result) as the gold standard. We posited that risk stratification with PHS would reflect the underlying incidence of PCa and therefore also affect the PPV of PSA testing.

In the Validation Set, 2,555 patients had positive PSA: 1,580 were then diagnosed with PCa, while 975 were designated controls without PCa. Because genotype information was collected in more cases than controls, we matched the overall ProtecT control:case ratio^10^ by taking a random sample of 471 cases with the 975 controls and calculating the positive predictive value of PSA testing without regard to PHS, as well as in subsets based on PHS percentile thresholds of <20^th^, >50^th^, >80^th^, and >95^th^. This process was repeated for a total of 1,000 random samples of 471 cases; mean and bootstrap estimate of the standard error were calculated for PPV in each PHS risk group.

To learn whether PHS impacts PPV within men of a given age category, we repeated the above PPV analysis for only Validation Set patients older than the median age of the group (60 years) and again for only those at or less than the median age.

The magnitude of PHS effect on expected age of onset was illustrated by calculating the PHS corresponding to percentiles among the young, healthy population within the Discovery Set: i.e., those controls with age <70 years. All percentiles reported in this manuscript refer to this population. Annualized incidence rate (*h_percentile_*) curves were calculated for each of various percentiles of the PHS from this population (1, 5, 20, 50, 80, 95, 99) with the median PHS (*PHS_median_*) is taken as baseline:

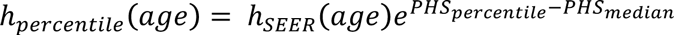

Application of an individualized PHS to screening decisions in the clinic would be facilitated by a readily interpretable translation of the PHS to terms familiar to the patient and physician. Thus, we introduce the “Prostate Cancer-Risk age” or PCaR age. An annualized incidence curve for the patient’s PHS is generated just as was done for the population percentiles above, which gives an estimate of PCa risk. Then, for example, if a 50-year-old man has a PCa risk equivalent to that of the general population at age 60, his PCaR age is 60.

A 95% confidence interval is calculated for PCaR age by estimating the variance due to both genotypes in the Discovery Set and the SNP parameter estimates from the PHS model as follows (details in Supplementary Methods).

The difference between PCaR age and true age (rounded to the nearest integer) is termed age. The PCaR age and age were calculated for every integer age between 40 and 95 years to assess whether age changed over time. This was done for all PHS percentiles listed above, as well as 0.1^th^ and 99.9^th^.

In a common clinical situation, a patient of a given age may present to his physician to discuss screening. To illustrate how PHS might influence this discussion, we identified the subset of Validation patients at approximately the median age, 60 years (57-63), to represent a typical patient. From this subset of 945 men around 60 years old, three groups were created: those whose PCaR age was also within the 57-63 interval, those with PCaR age <57, and those with PCaR age >63. We then calculated the PPV of PSA for these three groups using the same approach as before.

## Results

Of the 201,043 SNPs included in the data set, 2,415 were associated with increased risk of PCa in the trend test, with *p* <10^−6^. The stepwise regression framework then identified 54 of these SNPs that were incorporated into the Cox proportional hazards model (Supplementary Table S1). The 54 SNP parameter estimates (for the hazard of developing PCa) are combined with individual genotype to generate a polygenic hazard score. Kaplan-Meier curves indicate that the assumption of proportional hazards was reasonable in the final model (Figure 1).

**Figure 1:**
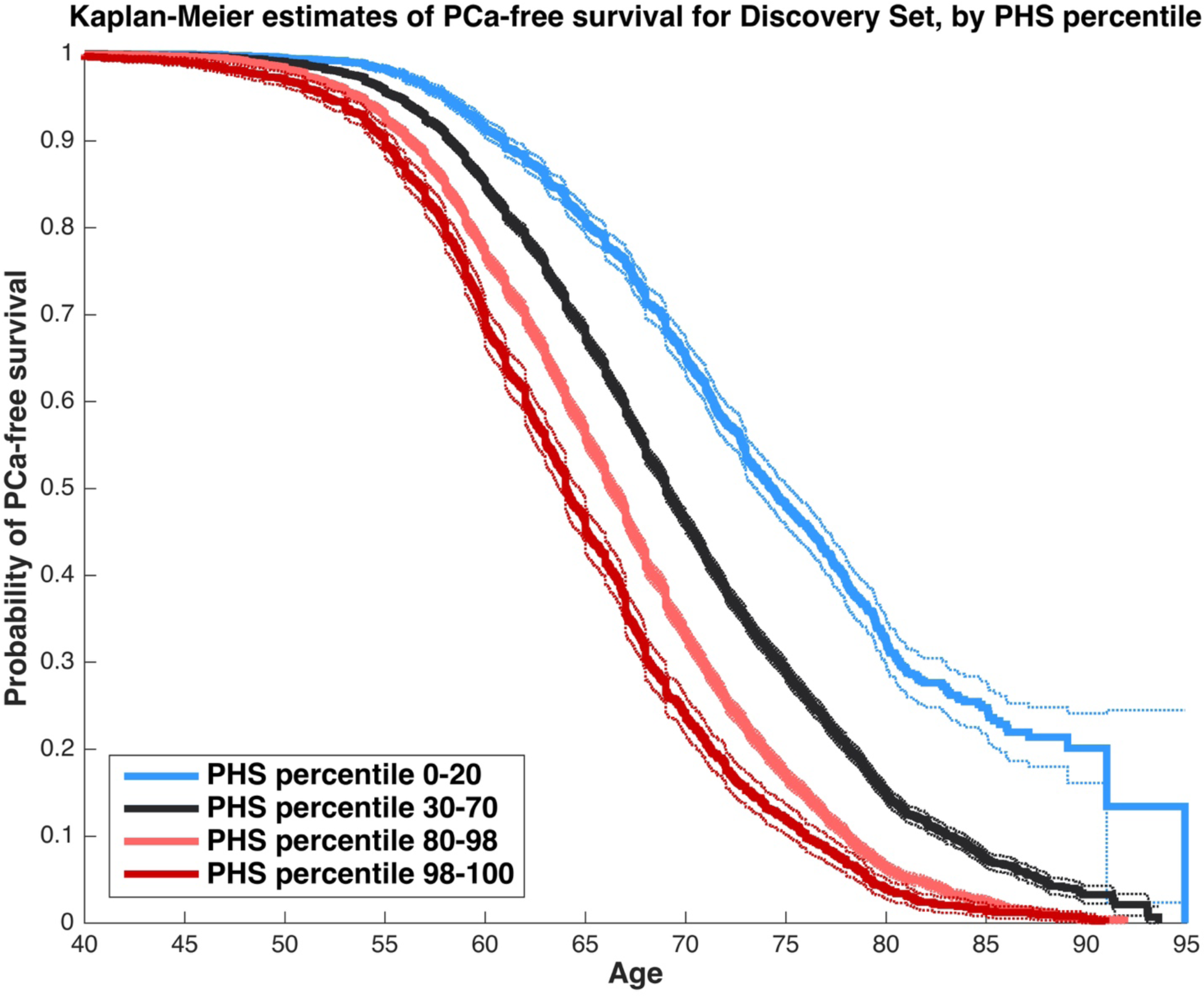
Kaplan-Meier estimates of prostate cancer-free survival for patients in the Discovery Set, grouped by PHS percentile ranges (as shown in the legend). PHS percentiles are in reference to the distribution of PHS within the 11,190 controls in the Discovery Set who were under 70 years old. Dotted lines are 95% confidence intervals for the corresponding curves. Time of “failure” is age at prostate cancer diagnosis. Controls were censored at age of observation. These curves demonstrate that the proportional hazards assumption holds for this PHS model.

In the independent Validation Set from the ProtecT study, a Cox proportional hazards model showed that PHS significantly predicted age of prostate cancer onset (*z* =15.4, *p* =10^−53^.

Positive predictive value of PSA testing in the Validation Set is plotted in Figure 2. PPV was lower among patients with a low PHS, and higher among patients with progressively higher PHS. Patients with PHS <20^th^ percentile had PPV 0.18, while those with PHS > 95^th^ percentile had PPV 0.52. Within the ≤60 and >60 age groups, PHS stratification still resulted in notable changes in the PPV of PSA testing (Figure 3).

**Figure 2:**
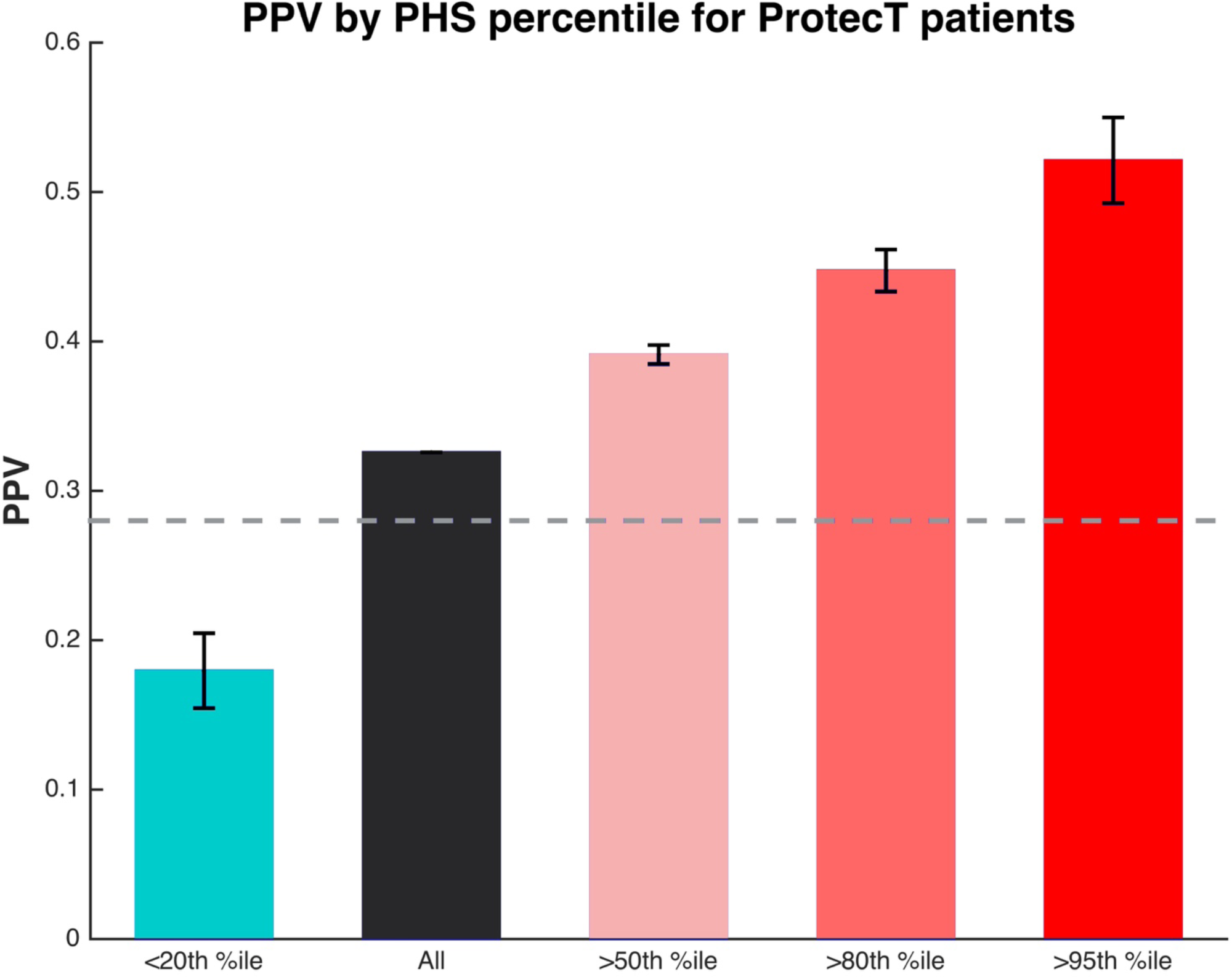
Positive predictive value (PPV) of PSA testing by PHS percentile thresholds for patients in the Validation Set. Percentiles refer to the PHS distribution among young controls in the Discovery Set. Error bars are the bootstrap estimate of the standard error for random samples of cases in the Validation Set (see Methods). For reference, the expected PPV for PSA testing at this threshold is displayed as a gray, dashed line, based on a pooled analysis^3^.

**Figure 3:**
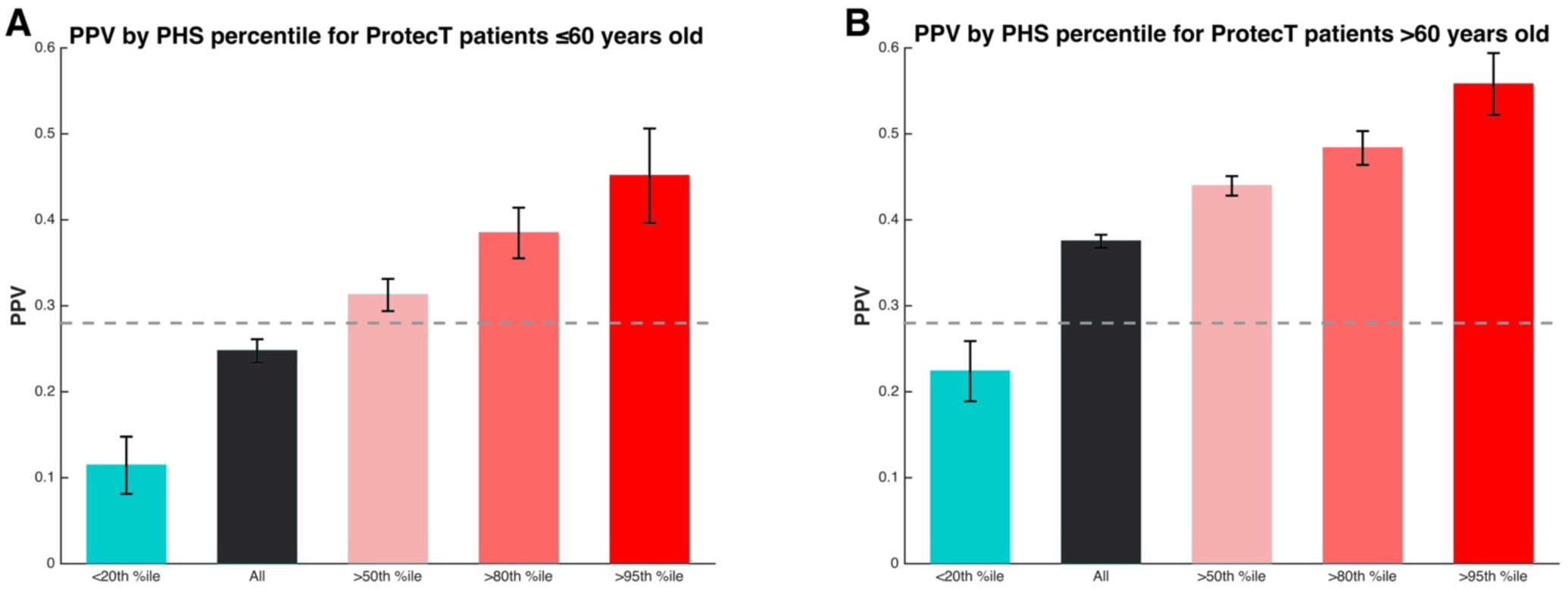
Positive predictive value (PPV) of PSA testing by PHS percentile thresholds for patients in the Validation Set, split by age group. (A) shows results for patients at or below the median age (60), and (B) shows results for patients older than the median age. For both panels: percentiles refer to the PHS distribution among young controls in the Discovery Set. Error bars are the bootstrap estimate of the standard error for random samples of cases in the Validation Set (see Methods). For reference, the expected PPV for PSA testing at this threshold is displayed as a gray, dashed line, based on a pooled analysis^3^.

Absolute risk of PCa for the general U.S. population was estimated with linear regression using data from the SEER database from 1995 to 2012^12,14–16^. The resulting model for hazard rate (*h_SEER_*) had *R^2^* =0.88 and *p* =10^−5^ (Figure 4):

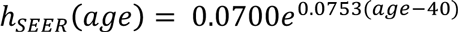

Annualized incidence and survival curves corresponding to PHS percentiles (among controls <70 years old) are shown in Figure 5. A table of prostate cancer-risk (PCaR) ages for various PHS levels demonstrates shows that the expected age of PCa onset is modulated by 20 years between the 1^st^ and 99^th^ PHS percentiles and by nearly 50 years between the 0.1^th^ and 99.9^th^ percentiles (Table 2).

**Figure 4:**
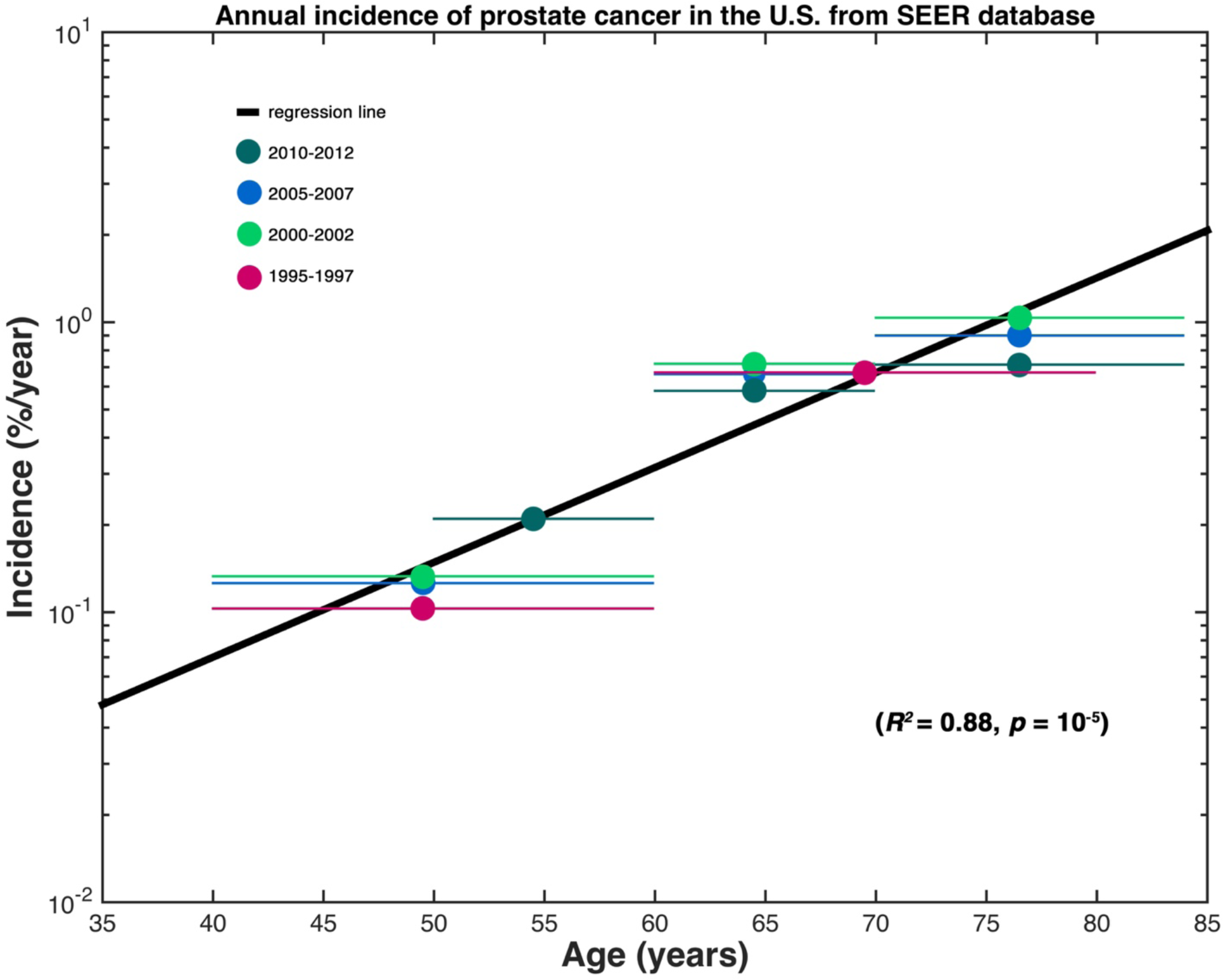
Dots represent the annual, age-specific incidence estimated from each age range (shown as horizontal lines of matching color) from U.S. population data in the SEER database. Dot color corresponds to the years the data were collected from, as shown in the legend. The black line is the result of linear regression for an exponential curve to give a continuous model of age-specific incidence in the U.S. population.

**Figure 5:**
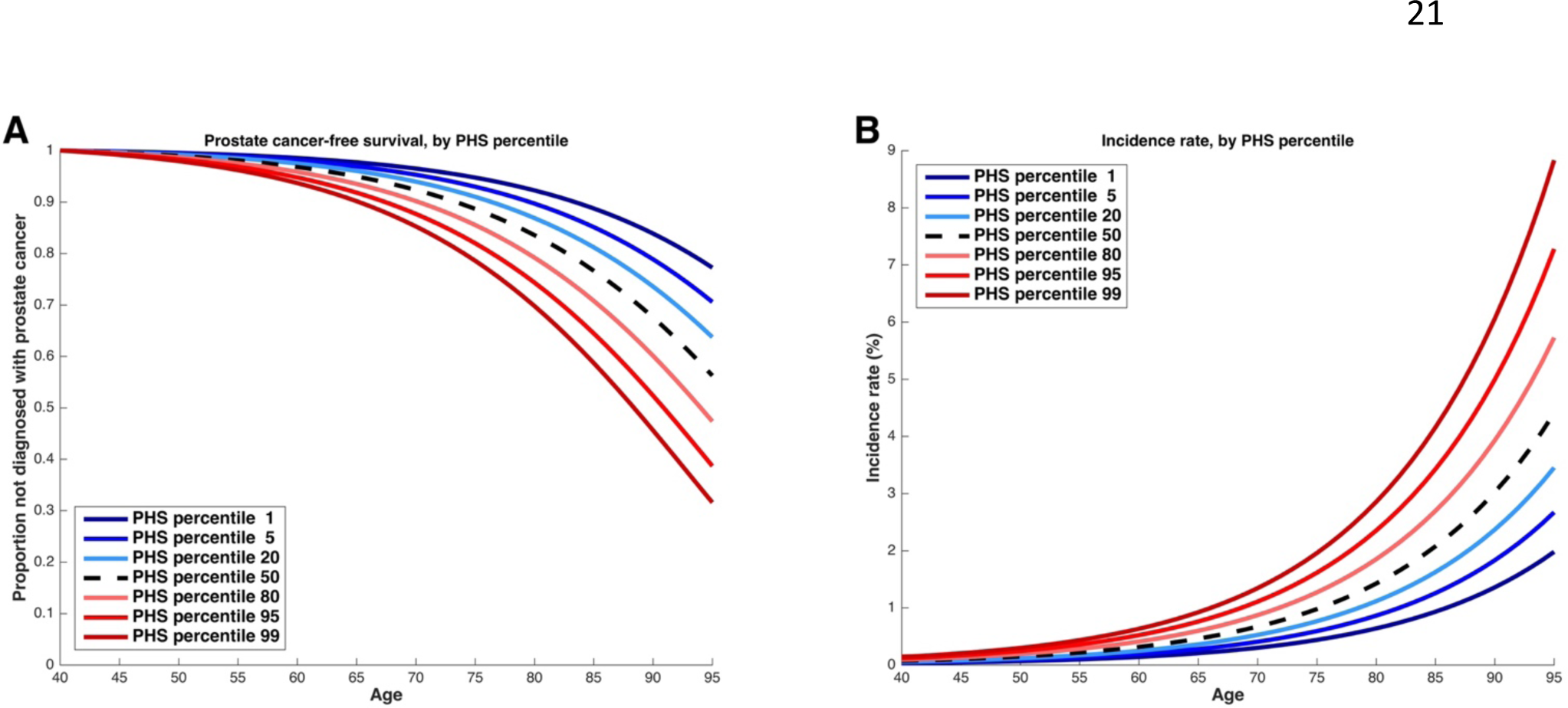
(A) Prostate cancer-free survival curves derived from PHS hazards, with U.S. population data taken as the median risk. PHS percentiles are in reference to the distribution of PHS within the 11,190 controls in the Discovery Set less than 70 years old. Blue lines represent genetic risk lower than the median, and red lines represent genetic risk higher than the median. (B) Incidence rate curves by age for the same risk levels as in (A).

**Table 2:**
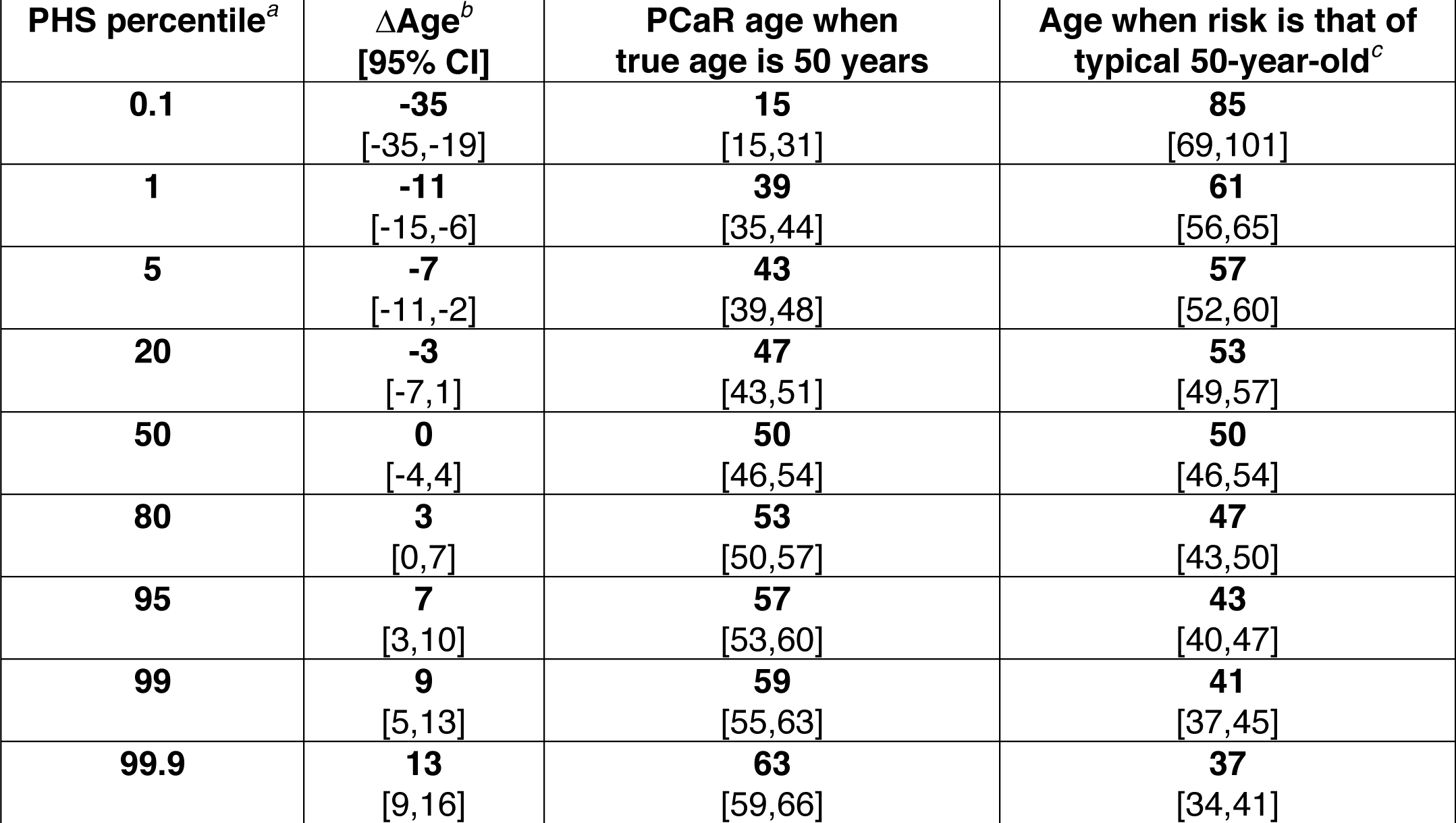
Prostate cancer-risk (PCaR) age

a PHS percentile among young (<70 years old) controls within Discovery Set

b ΔAge = PCaR age – true age

c Risk of typical 50-year-old defined as overall population incidence at age 50

Qualitatively, the curves in Figure 5 appear to maintain relatively consistent horizontal shifts relative to their neighbors over the age range studied. Quantitatively, this is confirmed by age, which remained the same for each PHS percentile across a true age range of 40 to 95. Thus, age was taken to be approximately constant for each PHS percentile and is reported in Table 2.

The PPV of PSA testing for Validation Set patients approximately 60 years of age (57-63) is shown in Figure 6. PPV was lower for those with PCaR age <57 and higher for those with PCaR age >63.

**Figure 6:**
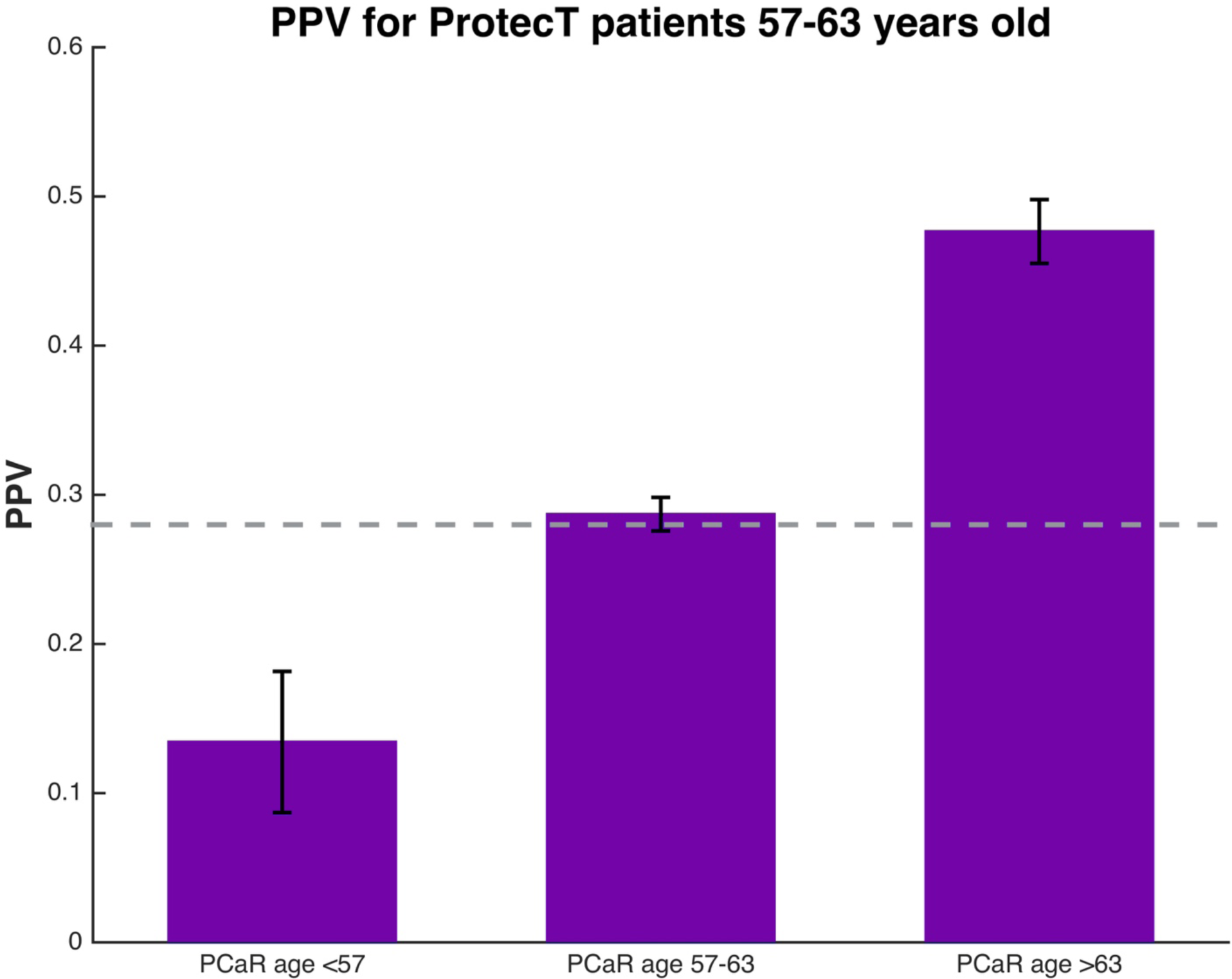
Application of Prostate Cancer-Risk (PCaR) age to the question of whether to screen a 60-year-old man. PCaR age is his true age adjusted by PHS level. For example, if his PCaR age is 71 years, his estimated risk matches that of a typical 71-year-old. Alternatively, a PCaR age of 52 would mean his PCa risk is similar to a typical 52-year-old. The bar plot shows results for all men from the Validation Set with age approximately 60 years (57-63), grouped by their calculated PCaR age: <57, 57-63, or >63. The positive predictive value (PPV) of PSA testing is shown for these groups. Error bars are the bootstrap estimate of the standard error. For reference, the expected PPV for PSA testing is displayed as a gray, dashed line, based on a pooled analysis^3^. ~68% of men had PCaR age 57-63 and PPV close to the expected population value. ~26% and ~7% of men were in the higher and lower groups, respectively.

## Discussion

Genetic information may be the ideal guide for deciding whether an individual patient needs prostate cancer screening^8^. The polygenic hazard score described here represents a personalized genetic assessment of a patient’s age-related prostate cancer risk that can inform both whether and when to order screening tests. In a survival analysis using patients from an independent clinical trial, PHS was a remarkably strong predictor (*p* =10^−53^) of age at PCa diagnosis. Furthermore, risk-stratification with PHS had considerable impact on the positive predictive value of PSA testing. For example, among patients with elevated PSA, only 18% of those with low PHS were true positives, whereas over half of those with high PHS had prostate cancer (Figure 3). As PHS is representative of a man’s fixed genetic risk, it can be calculated long before onset of PCa and substantially inform the decision of whether he should undergo PCa screening.

Because PCa incidence is highly dependent on age^12^, one must decide not only whether to screen but also at what age to consider it. Prostate Cancer-Risk (PCaR) age incorporates a patient’s true age and genetic risk to give an adjusted age that relates his current risk of PCa to that of the age-specific population average. For example, a physician who normally discusses the risks and benefits of PSA screening with her patients starting at age 50 could shift the timing of that conversation for each patient according to his PCaR age. Using the U.S. SEER database as the population average, we show here that PHS modulates PCa-free survival curves by 20 years between the 1^st^ and 99^th^ percentiles (Figure 5). The comparison is even more extreme for PHS percentiles 0.1 and 99.9 (Table 2), with some men not reaching the risk of a typical 50-year-old until age 85, while others reach that risk at age 37. With nearly 40 million men aged 50-70 years in the U.S. alone^18^, almost 80,000 would presumably fall into one of these extreme PHS categories.

To illustrate the usefulness of PCaR age, we consider a common clinical scenario where a man in clinic asks about screening. We assume his age is 60, the median for the ProtecT cohort. Figure 6 shows results for ProtecT patients approximately 60 (57-63) years old and suggests that if this 60-year-old man’s PCaR age is close to his true age (i.e., still 57-63), the PPV of a PSA test for him now is 29%, close to the average for PSA screening in general^3^. If his PCaR age is under 57, the PPV drops to around 13%, and he might be reassured that a PSA test is not necessary. On the other hand, if his PCaR age is over 63, the PPV is approaching 50%, which might make PSA more informative.

PSA screening is controversial, but most guidelines recommend individualized discussion between physicians and patients^12,14–16^. PHS affords a quantitative and understandable way to evaluate individual risk that could prove pivotal in these discussions. The results here suggest PHS can be used to identify a large percentage of men for whom forgoing or delaying PSA screening makes sense. At the same time, PHS can also identify men with a high risk of developing PCa at a young age and who therefore may benefit from early detection through PSA screening.

Another concern with PSA screening is overtreatment of indolent disease. Genetic prediction of aggressive PCa alone has proven elusive^19^, and the problem is compounded by the propensity of initially low-risk tumors to progress^20,21^. Datasets describing the initial tumor characteristics are not enough—tumors that develop aggressive features over time must also be identified if aggressive-PCa-only predictors are to be effective. Active surveillance is one answer to overtreatment that avoids up-front aggressive treatment but still allows intervention if the tumor progresses. Indeed, most tumors will eventually require treatment^21,22^, and earlier treatment prevents development of metastatic disease^22^. Hence, avoiding screening altogether in patients who may develop PCa at a young age does carry risk of considerable morbidity.

While PHS was applied here to PSA screening, PHS itself is not specific to PSA testing. Rather, the PHS is predictive of patients’ underlying risk of PCa at a given age and therefore relates to pre-test probability—and, by extension, positive predictive value— within any screening strategy that might be adopted.

Cost effectiveness is a prominent concern in all discussions of healthcare policy. PHS has the potential to improve screening efficiency while also reducing overall costs. PHS need only be calculated once and is valid for a lifetime. The genotyping chip assay can be run for costs in the range of that for single-gene testing (e.g., *BRCA* mutation), informs multiple diseases^23^, and a saliva sample suffices. PSA screening and subsequent biopsies could thus be limited to those men at higher risk, while delaying or forgoing screening in those whose genetic makeup confers a low risk.

Prior studies have used GWAS-associated polymorphisms to predict risk of PCa using a case/control design^24–26^. However, epidemiologic data show that PCa risk is not a simple dichotomy of cases and controls, but rather is highly dependent on increasing age. Therefore, we opted for a survival analysis approach optimized for genetic prediction of age of PCa onset. The PHS can then be used in clinical decisions, where age plays a critical role. If a man has a high risk of developing prostate cancer at age 95, this is a very different clinical situation from a man at high risk at age 55.

There are several limitations to this study. It is inherently a retrospective analysis, but the Discovery Set data come from large studies carried out in multiple institutions and nations; the Validation Set, too, comes from an independent, large, prospective trial. The absolute risk models shown in Figure 5 are only as accurate as the population data upon which they are based, which reflect *diagnosed* prostate cancer in the U.S. It is important to note, though, that PHS is a measure of hazard and therefore could be readily applied to other population incidence curves to estimate absolute risk in those populations. The age range of the Validation Set is limited to only 50-70 years; fortunately, this includes the age where screening is believed to have the most benefit^12,14–16^. Finally, race in this PHS model is limited to European ancestry. Validation of PHS in other racial groups—and, if necessary, custom models for each—is needed; our group plans to investigate this important question.

In conclusion, we describe here the development of a new polygenic hazard score for personalized genetic assessment of individual, age-associated prostate cancer risk. This score has been validated in an independent data set, demonstrating accurate prediction of prostate cancer onset. Moreover, PHS is shown to predict the utility of PSA testing for an individual patient and can guide the decision of whether and when to screen for prostate cancer.

## Author Contributions

TMS, CCF, VZ, DSK, IGM, OAA, and AMD designed the study. RAE, DFE, ZSKJ, AAAO, SBG, KM, HG, FW, MA, JS, CSi, TLJT, BGN, SFN, MW, RB, MAR, PI, TJK, RCT, DEN, JLD, FCH, PPh, NP, KTK, CM, WV, ML, KH, ASK, CC, DW, WK, LCA, HB, KC, KUS, JYP, TAS, CSl, RKan, VM, JB, JAC, AS, APCBR, MRT, PPa, SM, HP, AM, and AK collected the data. TMS, CCF, IGM, OAA, and AMD performed the literature search. TMS, CCF, YW, VZ, RKar, DSK, and AMD performed the data analysis. TMS, CCF, RKar, JKP, DSK, OAA, and AMD interpreted the data. TMS, RKar, JKP, DSK, OAA, and AMD created the figures. TMS, CCF, OAA, and AMD wrote the manuscript. All authors reviewed the manuscript, added appropriate revisions, agreed to submission for publication, and approved the final version.

## Research in Context

### Evidence before this study

Prostate cancer (PCa) screening with prostate-specific-antigen (PSA) testing can lead to early detection of PCa and allow for curative treatment, but universal screening also has considerable disadvantages for men who may never develop life-threatening disease. Whom to screen and at what age to do so remain unclear.

Genetic studies have shown that single-nucleotide polymorphisms (SNPs) have modest predictive value for PCa risk, but that a combination of genotype information from multiple SNPs can give a more robust PCa risk prediction. The practical, clinical utility of SNP-based prediction of expected age of PCa onset is not well understood.

### Added value of this study

This study presents and validates a novel polygenic hazard score that is an indicator of PCa-free survival. This polygenic hazard score (PHS) offers a relatively inexpensive assessment of an individual man’s *age-specific* PCa risk.

### Implications of all the available evidence

SNP-based polygenic hazard scores can provide objective, readily interpretable information to guide the decision of whether a given patient might benefit from PCa screening at each age in his life

